# A synthetic lethal screen for Snail-induced enzalutamide resistance identifies JAK/STAT signaling as a therapeutic vulnerability in prostate cancer

**DOI:** 10.1101/2022.11.15.516649

**Authors:** Kathryn E. Ware, Beatrice C. Thomas, Pelumi Olawuni, Maya U. Sheth, Nathan Hawkey, M Yeshwanth, Brian C. Miller, Katherine J. Vietor, Mohit Kumar Jolly, So Young Kim, Andrew J. Armstrong, Jason A. Somarelli

## Abstract

Despite substantial improvements in the treatment landscape of prostate cancer, the evolution of hormone therapy-resistant and metastatic prostate cancer remains a major cause of cancer-related death globally. The mainstay of treatment for advanced prostate cancer is targeting of androgen receptor signaling, including androgen deprivation therapy plus second-generation androgen receptor blockade (e.g., enzalutamide, apalutamide, darolutamide), and/or androgen synthesis inhibition (abiraterone). While these agents have significantly prolonged the lives of patients with advanced prostate cancer, the evolution of resistance to these treatments in nearly universal. This therapy resistance is mediated by diverse mechanisms, including both androgen receptor-dependent mechanisms, such as androgen receptor mutations, amplifications, alternatively spliced isoforms, and structural rearrangements, as well as non-androgen receptor-mediated mechanisms, such as lineage plasticity toward neuroendocrine-like or epithelial-mesenchymal transition (EMT)-like lineages. Our prior work identified the EMT transcriptional regulator Snail as critical to hormonal therapy resistance and commonly detected in human metastatic prostate cancer. In the current study, we sought to interrogate the actionable landscape of EMT-mediated hormone therapy-resistant prostate cancer to identify synthetic lethality and collateral sensitivity approaches to treating this aggressive disease state. Using a combination of high-throughput drug screens and multi-parameter phenotyping by confluence imaging, ATP production, and phenotypic plasticity reporters of EMT, we identified candidate synthetic lethalities to Snail-mediated EMT in prostate cancer. These analyses identified multiple actionable targets, such as XPO1, PI3K/mTOR, aurora kinases, c-MET, polo-like kinases, and JAK/STAT as synthetic lethalities in Snail+ prostate cancer. We validated these targets in a subsequent validation screen in an LNCaP-derived model of resistance to sequential androgen deprivation and enzalutamide. This follow-up screen provided validation of inhibitors of JAK/STAT and PI3K/mTOR as therapeutic vulnerabilities for Snail+ and enzalutamide-resistant prostate cancer.

## Introduction

The treatment landscape of prostate cancer exemplifies the “two truths” of cancer treatment [1]: While tremendous progress has been made to improve patient outcomes, there also remains an urgent need to overcome the significant challenges imposed by the evolution of treatment resistance and metastasis. From the groundbreaking studies of Huggins and Hodges [2] to the development of novel, second-generation androgen receptor inhibitors [3-8], and anti-androgens [9, 10], much of the existing treatment options for prostate cancer are currently focused on targeting the androgen receptor (AR) signaling axis. These agents have demonstrated significant clinical benefit; however, progression of men treated with these agents in the metastatic, castration-resistant setting is nearly universal.

The evolution of resistance to AR signaling inhibitors is mediated by heterogeneous genetic and non-genetic pathways that include both AR-dependent and AR-independent mechanisms (reviewed in [11]). Among these heterogeneous mechanisms, phenotypic plasticity is a central hallmark of AR signaling inhibitor resistance [12]. This phenotypic plasticity occurs along multiple, interconnected cellular lineage axes, such as stemness [13, 14], epithelial/mesenchymal [15-18], luminal/basal [19, 20], and neuroendocrine-like lineages or cell states [21, 22]. Phenotypic plasticity along these axes often leads to a loss of AR expression/activity and dependency [23], as well as additional aggressive features that promote survival and metastasis [24, 25]. New approaches are needed to capitalize on these emerging phenotypic states for therapeutic benefit.

Targeted therapy alters the ecological fitness landscapes of cancer in multiple ways [26]. The altered fitness landscape of the drugged environment can promote aggressive biology, but can also induce “collateral sensitivities” to novel agents [27]. This concept, also known as negative cross resistance, has been applied to identify new strategies to treat the evolution of resistance in bacterial infections [28], malaria [29], herbicides [30], and pesticides [31].

In the present study, we combined high-throughput screens with multiparameter endpoint measurements from transcription-based reporters, confluence, and cell viability assays to characterize the therapeutic landscapes of Snail-mediated EMT, enzalutamide resistance, and AR activity (**Fig. 1A**). Our analyses pinpoint histone deacetylases (HDAC), protein kinase A (PKA), PI3K/mTOR, and Janus Kinase (JAK) as key collateral sensitivities to Snail-mediated enzalutamide resistance in prostate cancer cells. Follow-up screens in a model of progressive adaptation to ADT and enzalutamide resistance verified the relevance of these pathways as novel therapeutic vulnerabilities for enzalutamide-resistant prostate cancer (**Fig. 1B**). These analyses provide a deeper understanding of the therapeutic vulnerabilities induced by epithelial plasticity and enzalutamide resistance.

**Figure 1.**
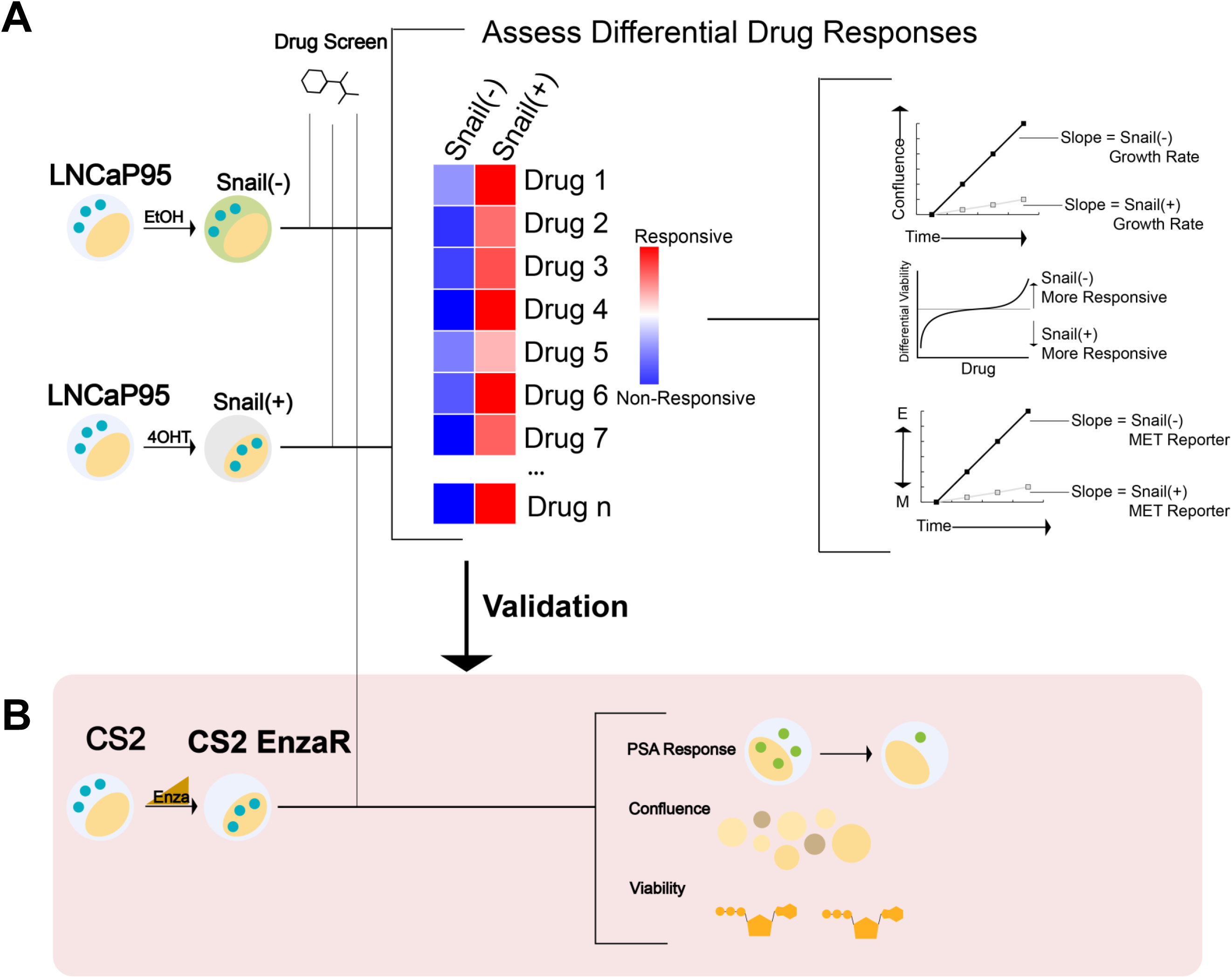
Workflow schematic for synthetic lethal and collateral sensitivity screens. **A**. A high-throughput screen was performed in LNCaP95-Snail cells to assess differential response across multiple endpoints of confluence, viability (CellTiter Glo), and EMT status via a fluorescence-based reporter. **B**. Screen schematic for a collateral sensitivity screen in enzalutamide-resistant CS2 cells. Endpoints included PSA reporter response, confluence, and viability (CellTiter Glo).

## Materials and Methods

### Cell culture models

LNCaP95-Snail and CS2 enzalutamide-resistant cells were cultured in RPMI containing 10% charcoal stripped Fetal Bovine Serum (Sigma) and 1% penicillin/streptomycin (Life Technologies). CS2 enzalutamide-resistant cell populations were maintained in the presence of 50 μM enzalutamide. Cell lines were maintained in standard tissue culture-treated plasticware within a humidified incubator at 37°C and 5% CO_2_. LNCaP95 cells stably expressing inducible Snail were generated as previously described [15]. Induction of Snail nuclear translocation was mediated by the addition of 4-hydroxy-tamoxifen (4OHT) at a concentration of 20 nM Ethanol (EtOH) was used as a vehicle control. All cells were authenticated by the Duke DNA Analysis Facility using analysis of short tandem repeats and were verified to be mycoplasma-free.

### Development and testing of MET and PSA reporter lines

We adapted the GIIIcI^2^ MET reporter [32, 33] for lentiviral transduction by cloning the previously-described vector into the lentiviral vector pLVX-puro using restriction enzymes EcoRI/SmaI. The PSA reporter was synthesized in the lentiviral expression plasmid, pLV[Exp]-Puro by VectorBuilder to include 2 Kb of the proximal PSA promoter upstream of the EGFP open reading frame. Cells stably expressing inducible Snail (Addgene plasmid #18798) or indicated reporter plasmids were generated by transduction of LNCaP95 or CS2 cells as described:

https://www.addgene.org/protocols/generating-stable-cell-lines/. Confluence and fluorescence were measured with and without EMT induction using Snail activation as described above. For PSA-GFP expressing cells, confluence and fluorescence was quantified with and without AR activation using synthetic androgen R1881 at 1 nM.

### High-throughput drug screening

High-throughput screens were performed in collaboration with the Duke Functional Genomics Shared Resource as previously described [34-37]. Briefly, compounds from the Bioactives library (SelleckChem) were stamped in triplicate into 384-well plates at a final concentration of 1 μM using an Echo Acoustic Dispenser (Labcyte, Indianapolis, IN, USA). Cells and media were subsequently dispensed into plates using a WellMate (Thermo Fisher, Waltham, MA, USA) at a density of 2,000 cells/well for each cell line. Confluence was quantified using an IncuCyte S3 live cell imaging system. GIIIcI^2^ and PSA-GFP readouts were quantified by IncuCyte imaging at 24, 48, 72, and 96 hours. CellTiter Glo was added at 96 hours, and luminescence was read using a Clariostar plate reader (BMG, Berlin, Germany).

### RNA-Seq analysis of EMT scores

Quantification of EMT status for each sample was performed using the following three independent methods: 76GS, KS, MLR, each of which uses a unique algorithm and gene set. The 76GS scores were calculated based on the expression of 76 genes [38]. Higher scores correspond to more epithelial states. A 76GS score > 0 typically indicates an epithelial phenotype and < 0 indicates a mesenchymal phenotype. The score for each sample is computed as the weighted sum of expression values of 76 genes, with the weight factor being the correlation of expression values of that gene with that of CDH1 in the given dataset. KS score was determined based on a Kolmogorov–Smirnov two-samples test [39]. Using a 218 gene signature, the cumulative distribution functions are estimated for mesenchymal and epithelial signatures, and the maximum difference in cumulative distribution functions is retained as the statistic for the two sample-KS test. KS score ranges from [–1, 1], with negative and positive scores representing mesenchymal and epithelial phenotypes, respectively. MLR scores are provided on a scale of [0, 2]; higher scores are associated with more mesenchymal samples. Using an ordinal multinomial logistic regression, the score encompasses an order structure, with a hybrid epithelial/mesenchymal signature situated between the epithelial and mesenchymal phenotypes. Scores are calculated based on the probability assigned for each sample to belong to one of the three phenotypes.

### Data analysis

The primary objective for the high-throughput screen of LNCaP95-Snail cells was to identify synthetic lethality for Snail+ cells. Snail-cells (EtOH-treated vehicle controls) were used as a reference control to calculate differential effects across all parameters. The primary objective for the high-throughput screen of CS2 enzalutamide-resistant cells was to identify collateral sensitivities for enzalutamide-resistant cells. The central hypothesis for this work was that activation of key pathways in Snail+, enzalutamide-resistant prostate cancer can be exploited for therapeutic benefit through synthetic lethal and collateral sensitivity approaches. Experimental data were visualized and analyzed in GraphPad Prism 9. Analysis of cell viability by CellTiter Glo was performed by normalizing to the average of all empty (non-drug) wells. Imaging of confluence and GFP was compared using repeated measures ANOVA. Linear regression was used to assess correlations between screen analysis parameters, and outliers were considered to fall outside the 95% confidence interval bands. P-values <0.05 were considered statistically reliable.

## Results

### Fluorescence-based reporters enable real-time monitoring of epithelial plasticity

Prior studies have pinpointed the epithelial plasticity regulator, Snail, as both upregulated during AR inhibition [17] and a mediator of enzalutamide resistance through sustained androgen receptor signaling [15]. In the present work we sought to develop a system to identify novel collateral sensitivities to Snail-induced resistance to enzalutamide. To do this we turned to a Snail inducible LNCaP95 cell line system in which Snail is fused to an estrogen receptor mutant (ER^mut^) in which 4-hydroxy-tamoxifen (4OHT) acts as an agonist (**Fig. 2A**). Addition of 4OHT induces estrogen receptor-Snail fusion nuclear localization and activation of a Snail-mediated transcriptional program (**Fig. 2A**). Addition of 4OHT in the Snail-inducible LNCaP95 prostate cancer cell line leads to cell scattering, loss of cell-cell E-cadherin, and upregulation of the mesenchymal marker, vimentin (**Fig. 2B**). To track dynamics of Snail-mediated epithelial plasticity we adapted the GIIIcI^2^ fluorescence-based reporter [33] for lentiviral transduction. The GIIIcI^2^ reporter utilizes the lineage-specific alternative splicing within the ligand binding domain of FGFR2 to control EGFP expression based on epithelial or mesenchymal phenotype [33]. The EGFP open reading frame is interrupted by the FGFR2-IIIc exon and flanking introns (**Fig. 2C**). Splicing of FGFR2-IIIc in epithelial cells leads to fusion of the EGFP reading frame and subsequent EGFP expression while inclusion of the IIIc exon interrupts the EGFP reading frame and prevents EGFP expression (**Fig. 2C**). Treatment of LNCaP95-Snail cells with 4OHT leads to a reduction in confluence, consistent with the known relationship between Snail and cell cycle arrest [40] (**Fig. 2D**). Similarly, Snail induction also induces robust inhibition of EGFP expression (**Fig. 2E**) consistent with inclusion of the mesenchymal FGFR2-IIIc exon. A loss of EGFP signal in Snail+ cells is also evident by fluorescence imaging of Snail-(EtOH) and Snail+ (4OHT) cells (**Fig. 2F**). EGFP expression from the GIIIcI^2^ reporter is also consistent with endogenous FGFR2 splicing, in which 4OHT induces a switch from the IIIb to IIIc isoforms, as observed by isoform-specific restriction digestion of FGFR2 RT-PCR products (**Fig. 2G**).

**Figure 2.**
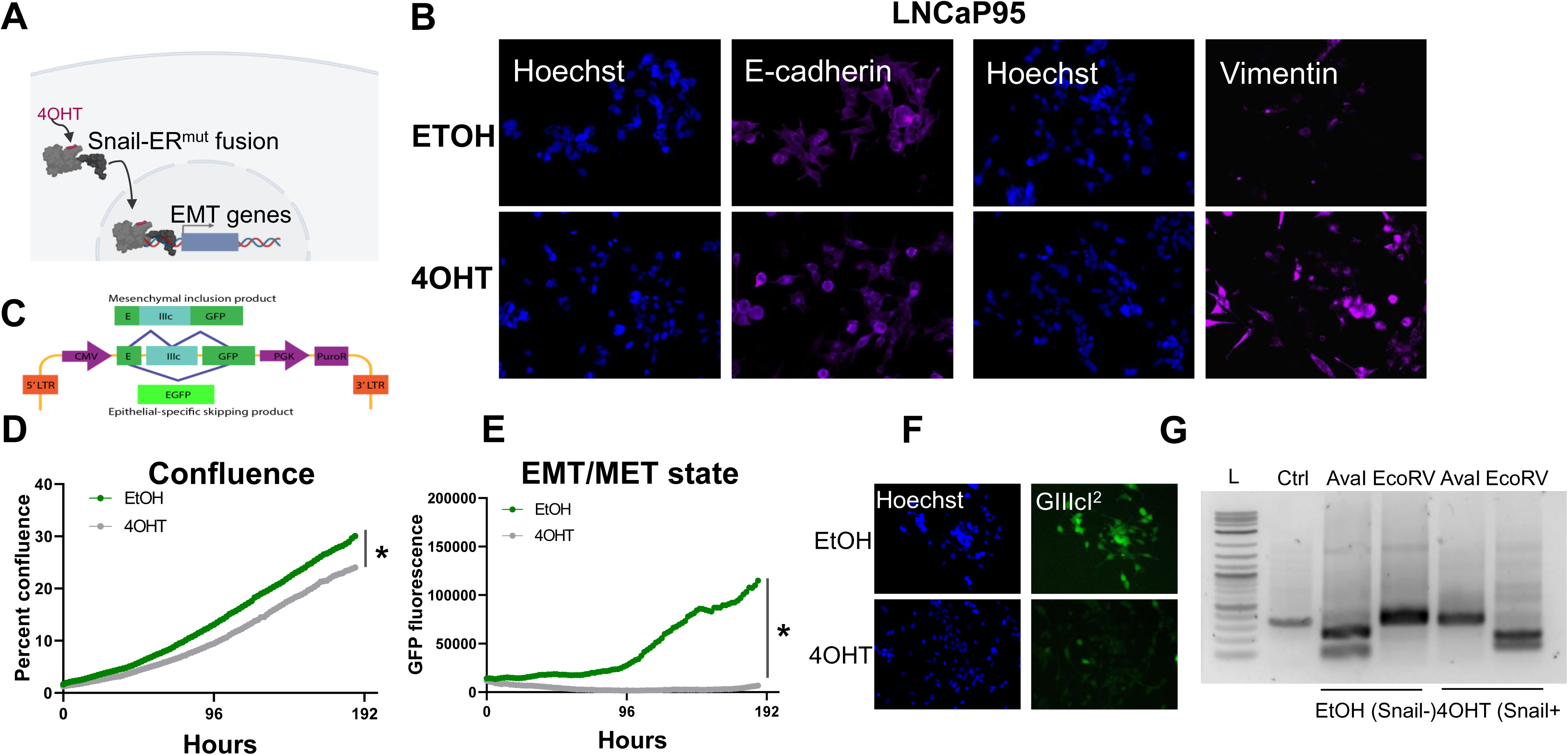
Fluorescence-based reporters to visualize EMT dynamics in a Snail-inducible model. **A**. Schematic illustration of a Snail-inducible model. **B**. Immunofluorescence staining of LNCaP95-Snail cells. EtOH serves as a vehicle for Snail induction. 4OHT induces localization of Snail and concomitant downregulation of E-cadherin and upregulation of vimentin. **C**. Schematic of the GIIIcI^2^ EMT/MET alternative splicing reporter. **D**. IncuCyte imaging for LNCaP95-Snail confluence and **E**. EMT induction dynamics (GFP fluorescence). * = p<0.05. **F**. Fluorescence imaging of LNCaP95-Snail cells treated with EtOH or 4OHT for nuclear staining (Hoechst) and the GIIIcI^2^ EMT/MET reporter (green). **G**. Endogenous FGFR2 splicing analysis for Snail- and Snail+ LNCaP95 cells. L = 1Kb ladder, Ctrl = undigested PCR product; AvaI = FGFR2-IIIb-specific restriction digestion; EcoRV = FGFR2-IIIc-specific restriction digestion.

### High-throughput screens identify synthetic lethality to Snail-induced epithelial plasticity

We applied this Snail-inducible plasticity reporter system to identify compounds with synthetic lethality for Snail+ prostate cancer that could be subsequently validated for activity in models of enzalutamide resistance given the association between Snail expression and enzalutamide resistance [15]. To do this, we performed a high-throughput small molecule screen using the SelleckChem Bioactives compound library. The Bioactives library contains 2,100 small molecules annotated by target and pathway. The library was designed to include compounds that are structurally diverse, medicinally active, and cell permeable, including both FDA-approved and non-approved compounds [34, 36, 37]. Screen results were analyzed for cell viability/ATP production by CellTiter Glo at the four-day endpoint, and for cell growth rate and epithelial plasticity status by daily IncuCyte imaging of confluence and GIIIcI^2^ EGFP levels, respectively, for four days (**Fig. 3A**; **Supplemental Table 1**). Analysis of CellTiter Glo values for empty wells revealed a significant reduction in growth for Snail+ cells (**Supplementary Figure 1A**), which is consistent with the known role of Snail as a mediator of cell cycle arrest. Across the entire compound library 3.8% of compounds inhibited CellTiter Glo signal for Snail-cells of 50% or more, while 22% of the library inhibited Snail+ cells 50% or more (**Supplemental Table 1**). To identify compounds with differential sensitivity based on Snail expression, we analyzed the differential sensitivity of Snail- and Snail+ cells to all compounds in the library, with a 1.0 representing no difference in sensitivity. Drugs with values <1.0 differentially inhibit CellTiter Glo output of Snail+ cells while drugs with values>1.0 differentially inhibit CellTiter Glo output in Snail-cells (**Fig. 3B**).

**Figure 3.**
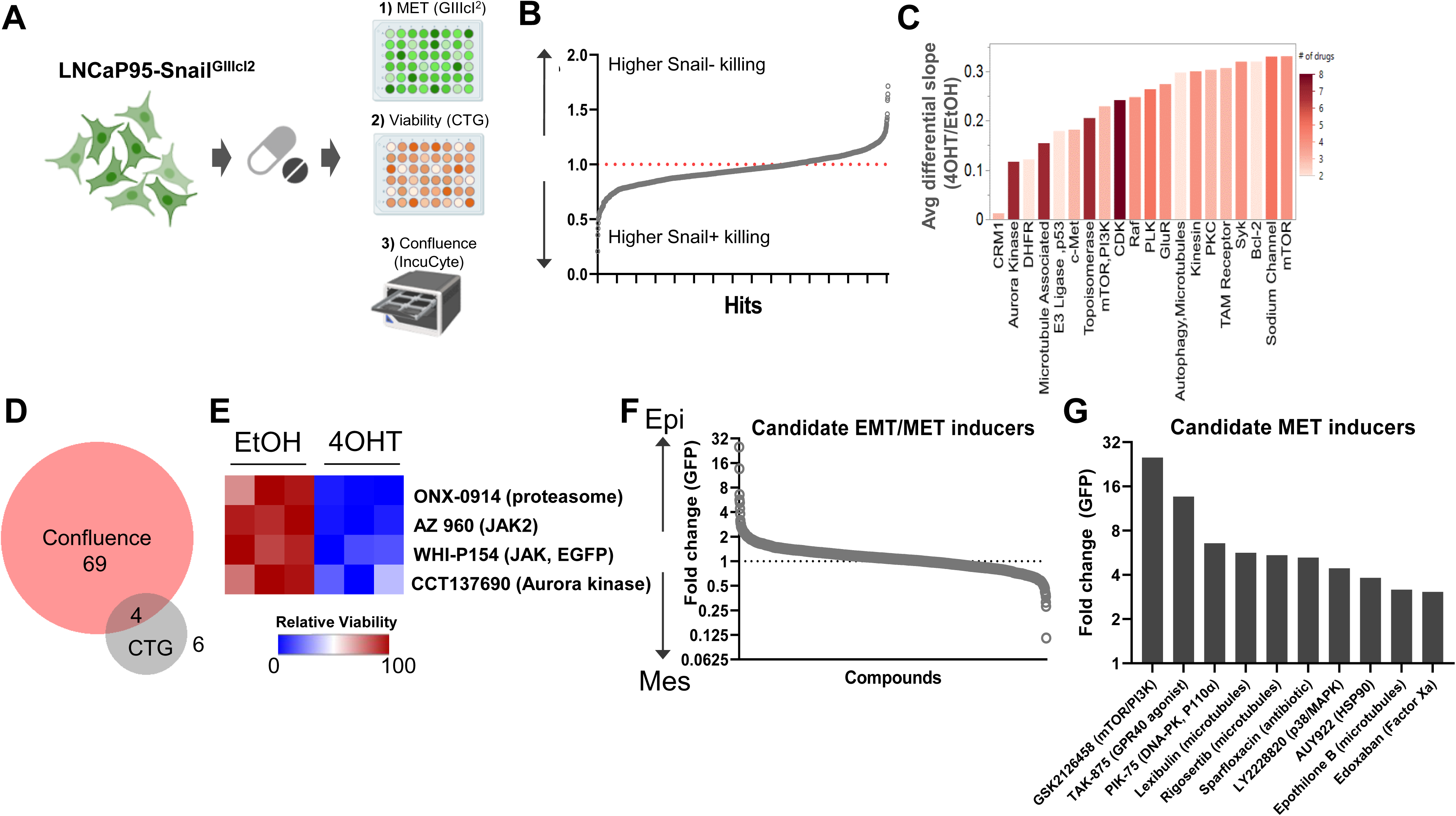
A synthetic lethality screen pinpoints potential therapies for Snail+ prostate cancer. **A**. Schematic of multi-assay screening strategy. **B**. Top hits with differential response in Snail – and Snail + cells. Below the 1.0 line indicates drug differentially inhibits Snail+ cells; above the line indicates drug differentially inhibits Snail-cells. **C**. Representative growth slope differences for top candidate agents with differential effects on Snail- and Snail+ cells. **D**. Top hits grouped by target/pathway ranked by differential slope; color indicates number of drugs per pathway. **E**. Venn diagram of overlap in compounds that altered both confluence and CellTiter Glo (CTG). **F**. Overlapping drugs with differential sensitivity in Snail+ cells for both confluence and CTG assays. **G**. Candidate EMT/MET inducers ranked by GIIIcI^2^ induction (higher GFP = more epithelial; lower GFP = more mesenchymal). **H**. Top 10 candidate MET inducing compounds, as estimated by EGFP expression from the GIIIcI^2^ reporter.

In parallel to CellTiter Glo, we also quantified differences in growth rate for all screen compounds with and without Snail induction. Cell confluence was moderately, but significantly, correlated with CellTiter Glo values when comparing all treatment conditions (**Supplemental Fig. 3B**). To identify collateral sensitivities based on growth rate we first calculated differences in slope of the growth rates between Snail-(EtOH) and Snail+ (4OHT) cells. This analysis is shown for a subset of compounds in **Fig. 1C**, with compounds in gray having little to no effect on cell growth of Snail-(EtOH) cells and these same compounds inhibiting growth in Snail+ (4OHT) cells. Subsequent annotation by target enabled identification of targets for which >2 drugs hit the same target. Top hits were ranked by their differential slope when comparing Snail+ to Snail-cells. Among these hits were inhibitors targeting signaling molecules and pathways known to be involved in lineage plasticity and prostate cancer therapy resistance, such as aurora kinase, c-MET, and mTOR/PI3K (**Fig. 3D**). Other targets included inhibitors of CRM1 (XPO1), a nuclear shuttling protein, cyclin-dependent kinases, polo-like kinases, and protein kinase C (**Fig. 3D**). To identify synthetic lethality for Snail+ cells, we focused on agents with <50% killing in Snail-(EtOH) cells and >50% killing in Snail+ cells by CellTiter Glo. Among these compounds, comparison of drugs that inhibited both CellTiter Glo production and growth rate by greater than 2-fold in Snail+ cells as compared to Snail-cells revealed four candidate compounds (**Fig. 3E**), including ONX-0914 (immunoproteasome inhibitor), AZ-960 (JAK2 inhibitor), WHI-P154 (JAK3 and EGFR inhibitor), and CCT137690 (aurora kinase inhibitor) (**Fig. 3F**).

We next attempted to identify compounds and pathways that inhibit Snail-induced EMT. To do this we first calculated the fold change in EGFP expression for each compound at day 4 as compared to day 1. The fold change in EGFP expression for 4OHT (Snail+) cells was divided by EtOH (Snail-) cells for each compound to identify drugs that were capable of overcoming Snail-mediated EMT. To ensure the gain in EGFP expression was not simply a function of cell growth inhibition or cell death, we compared the EGFP expression to the differential confluence in 4OHT-treated versus EtOH-treated cells. This analysis revealed a subset of compounds that led to differential re-activation of EGFP expression from the GIIIcI^2^ EMT/MET reporter while maintaining at least 50% viability or greater (**Fig. 3G**). These agents included GSK2126458 (mTOR/PI3K), three microtubule associated drugs, TAK-875 (GPR40 agonist), PIK-75 (DNA-PK, p110α), Sparfloxacin (antibiotic), LY2228820 (p38/MAPK), AUY922 (HSP90), and Edoxaban (Factor Xa) (**Fig. 3H**).

### The chemical landscape of collateral sensitivity to enzalutamide-resistant prostate cancer

Given the association between Snail-mediated EMT and enzalutamide resistance, we hypothesized that the evolution of enzalutamide resistance may also enrich for this EMT-like plasticity. To better understand these relationships between phenotypic plasticity and enzalutamide resistance we applied a series of EMT scoring metrics [41-43] to analyze RNA-Seq data from four independent pairs of enzalutamide-sensitive and enzalutamide-resistant cell line models [16]. Consistent with our hypothesis, enzalutamide-resistant cells exhibited a significant shift in scores toward a more mesenchymal-like gene expression signature (**Fig. 4A**). These overall trends were consistent across scoring metrics, with some exceptions for specific cell line pairs, depending on the scoring metric used (**Supplemental Fig. 2A, B**). Also consistent with this, treatment of LNCaP95-Snail(-) cells with enzalutamide led to an increase in nuclear localization of Snail (**Fig. 4B**). The enzalutamide-treated LNCaP95-Snail cells mirrored induction of Snail nuclear localization with 4OHT treatment (**Fig. 4C,D**). These analyses indicate that, compared to enzalutamide-sensitive cells, enzalutamide-resistant cells exhibit a more EMT-like phenotype.

To further extend the analysis of Snail-specific synthetic lethality, we next attempted to identify potential collateral sensitivities to this EMT-like enzalutamide-resistant phenotype. In order to accomplish this we performed a separate high-throughput compound screen on enzalutamide-resistant CS2 cells. The CS2 model is an LNCaP-derived subclone that was generated from long-term exposure to androgen deprivation through chronic culture in media containing charcoal-stripped fetal bovine serum [16]. Subsequent exposure of enzalutamide-sensitive CS2 cells to increasing doses of enzalutamide over approximately 6 months led to the development of an enzalutamide-resistant CS2 cell line model [16]. The CS2 enzalutamide-resistant model was transduced with a lentiviral PSA reporter in which the proximal promoter of PSA harboring androgen responsive elements is inserted upstream of the GFP reading frame (**Fig. 5A**). These CS2^PSA-GFP^ enzalutamide-resistant cells were screened using the Bioactives library to interrogate AR signaling (GFP), cell viability (CellTiter Glo), and cell growth (IncuCyte imaging) (**Fig. 5A**). To ensure the PSA reporter is responsive to androgen receptor signaling, cells were treated with the anabolic-androgenic steroid derivative, R1881, or enzalutamide. R1881 treatment led to a significant increase in GFP signal while enzalutamide had no effect on GFP expression in the enzalutamide-resistant CS2 model (**Fig. 5B**). The increase in GFP during R1881 treatment was not due to a change in confluence, as these treatments did not significantly alter cell confluence (**Fig. 5C**). Analysis of cell growth inhibition for the Bioactives screen at the pathway level in the CS2 enzalutamide-resistant cells pinpointed candidate collateral sensitivities of interest, including DNA-PK, cyclin-dependent kinases, histone deacetylases, PI3K, mTOR, CRM1, and PLK (**Fig. 5D**). Analysis of PSA reporter expression as a function of cell viability also revealed compounds targeting multiple receptors (androgen receptor, estrogen receptor, glucocorticoid receptor, dexamethasone) as inducers of PSA reporter activity (**Fig. 5E**) and compounds that target epigenetic modifiers as repressors of PSA reporter activity (**Fig. 5F**).

**Figure 4.**
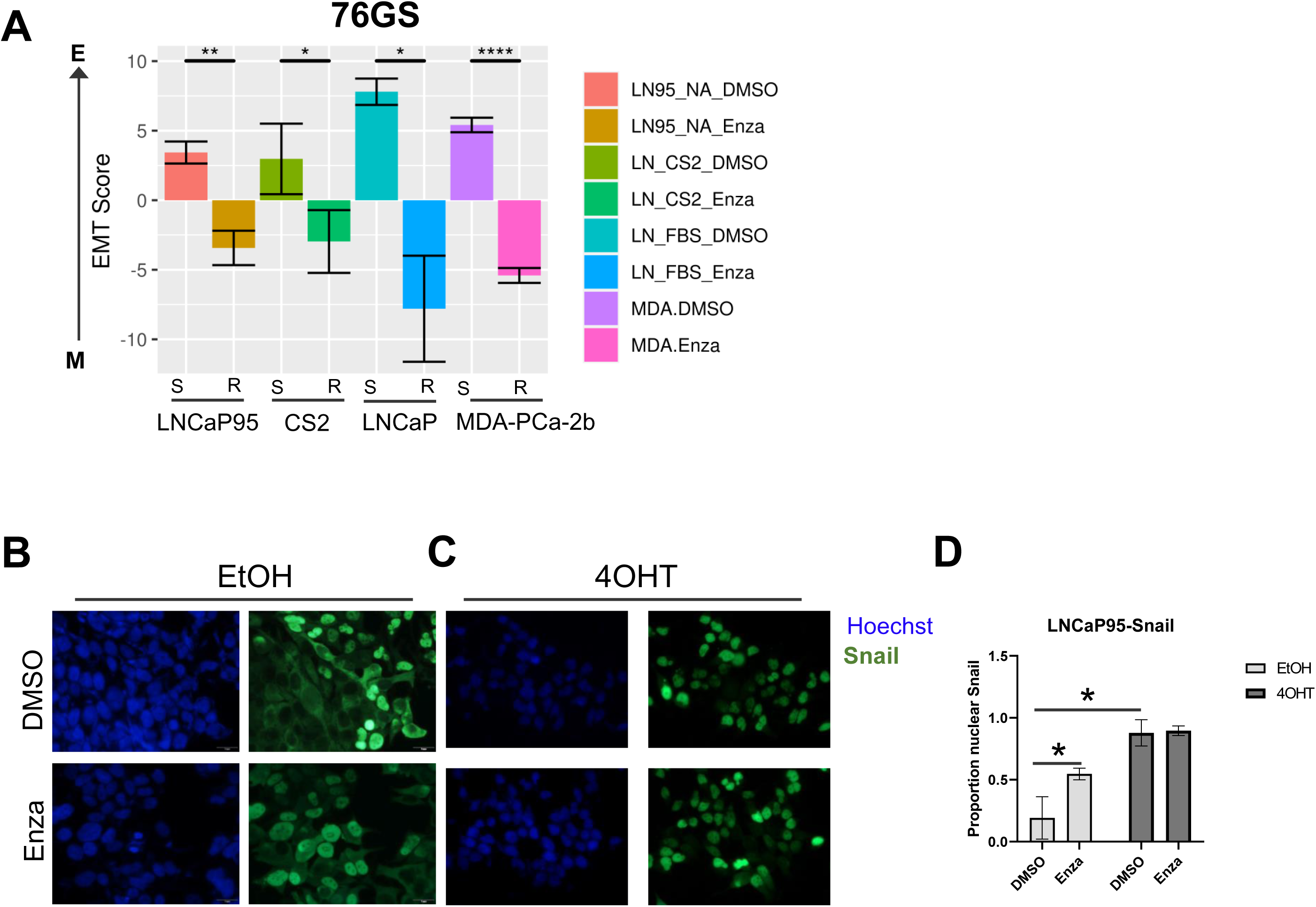
Enzalutamide induces epithelial plasticity. **A**. Analysis of EMT scores across three isogenic pairs of independently-derived enzalutamide-sensitive and enzalutamide-resistant cell line models using the 76GS EMT scoring metric; s = enza-sensitive; r = enza-resistant. **B**. Immunofluorescence staining of cell nuclei by Hoechst (blue) and Snail (green) in LNCaP95-Snail cells treated with EtOH (vehicle) and **C**. 4OHT (nuclear Snail) in the presence of vehicle or enzalutamide. D. Quantification of immunofluorescence by ImageJ. *=p<0.05

**Figure 5.**
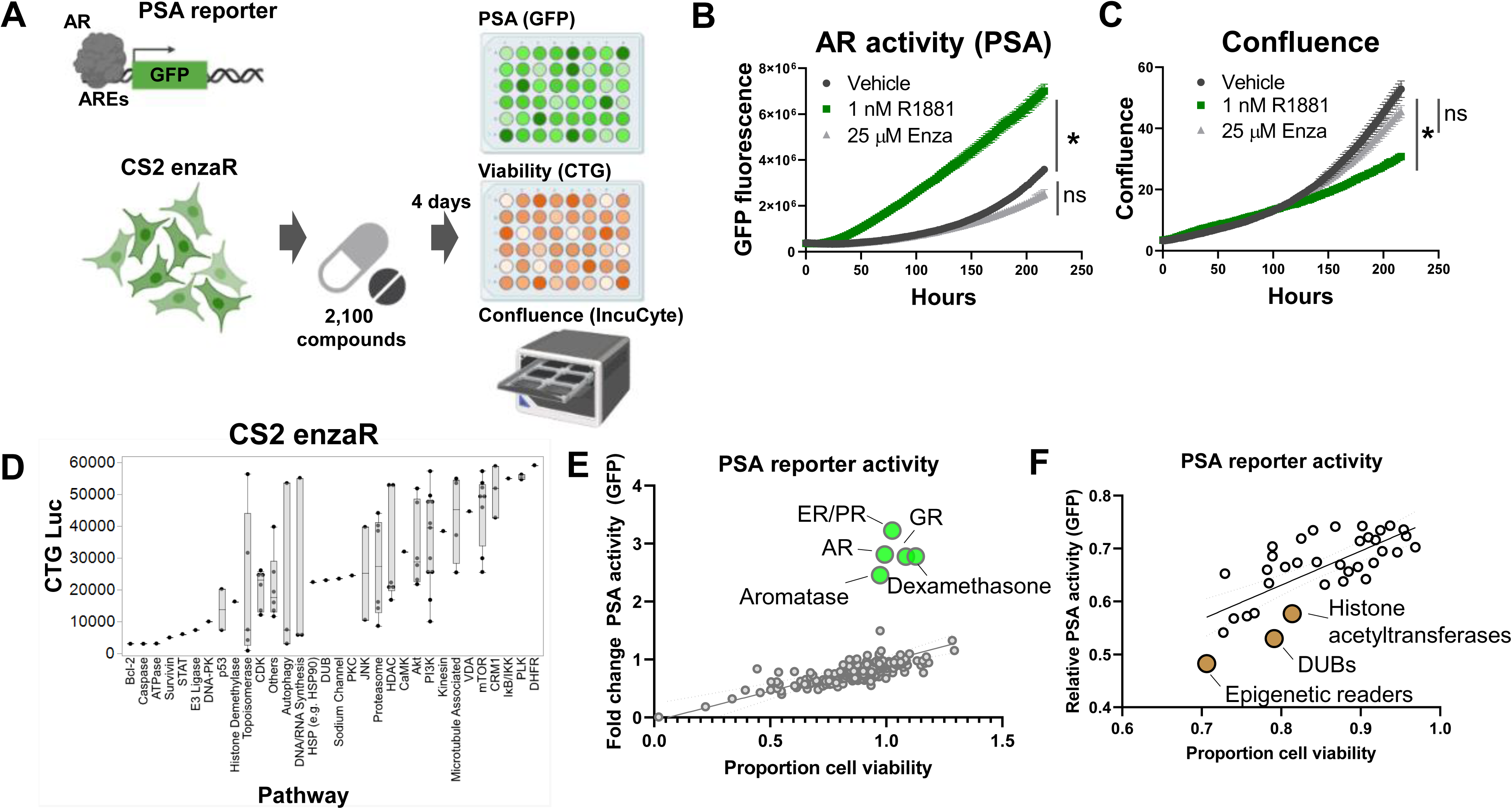
Collateral sensitivity screens identify candidate actionable pathways to treat enzalutamide-resistant prostate cancer. **A**. PSA reporter schematic and screening strategy. **B**. Validation of the PSA-GFP reporter system. **C**. Confluence quantification in CS2 enzalutamide-resistant model following exposure to R1881 and enzalutamide. **D**. Pathway-level analysis of top inhibitors targeting CS2 enzalutamide-resistant cells. **E**. Activators of PSA reporter activity (green dots); top candidates are labeled by pathway or with drug name. **F**. Inhibitors of PSA reporter activity (brown dots); top candidates are labeled by pathway.

To provide further validation of candidates, we plotted the relative cell viability by CellTiter Glo for compounds in the CS2 enzaR screen by cell viability (CellTiter Glo) in the LNCaP95-Snail screen (**Fig. 6A**). This analysis revealed a subset of drugs active in both screens. We ranked these top hits by a sum rank statistic that includes the rank of cell death by CellTiter for both screens as well as the differential confluence for Snail-vs. Snail+ cells (**Fig. 6B**). Top targets from this analysis including PI3K, mTOR, and the proteasome (**Fig. 6B**). Among this subset, AZ 960 (JAK2 inhibitor) and BGT226 (dual PI3K/mTOR inhibitor) were the most effective at inhibiting Snail+ cell confluence (**Fig. 6C, D**). Consistent with our observations of sensitivity to JAK2 inhibition, analysis of phospho-proteomics data from three previously-characterized pairs of enzalutamide-resistant lines [16], including CS2 enzalutamide-sensitive and -resistant lines demonstrates increased phosphorylation of STAT1, STAT2, JAK1, and JAK2 (**Fig. 6E-H)**, further highlighting the JAK/STAT signaling axis as a potential therapeutic vulnerability for Snail+ and enzalutamide-resistant prostate cancer.

**Figure 6.**
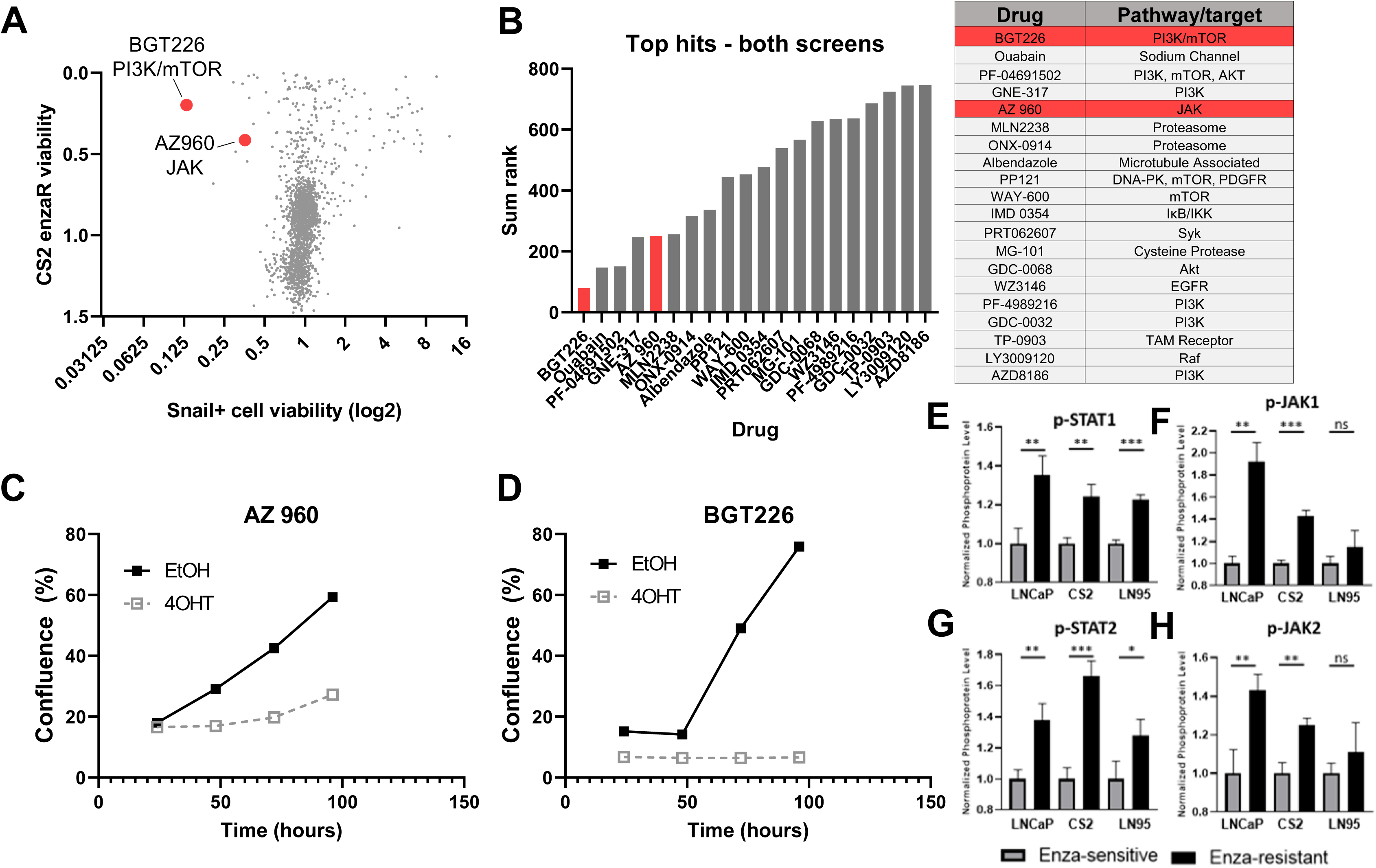
Comparison of candidate therapies for enzalutamide-resistant and Snail+ prostate cancer. **A**. Comparison of CS2 enzaR and Snail drug screen hits. **B**. Top hits for both screens based on a sum rank statistic that includes (CS2 enzaR confluence, Snail+ differential confluence, and Snail+ differential slope of growth rate). **C**. Growth curves for EtOH (Snail-) and 4OHT (Snail+) cells treated with AZ 960 (JAK inhibitor); and **D**. BGT226 (PI3K/mTOR inhibitor). **E**. Quantification of phospho-protein array data for p-STAT1, **F**. p-JAK1, **G**. p-STAT2, and **H**. p-JAK2 in three pairs of enzalutamide-sensitive and enzalutamide-resistant models (Ware et al. *Biorxiv*).

## Discussion

In the present study we sought to characterize the therapeutic vulnerabilities for enzalutamide-resistant prostate cancer. To do this we combined high-throughput small molecule screens with real-time imaging and endpoint assays to reveal chemical landscapes of synthetic lethality for Snail-mediated EMT and collateral sensitivities for enzalutamide-resistant cells. These screens identified multiple therapeutic vulnerabilities of Snail+ prostate cancer cells, including several with known functions in prostate cancer and/or EMT, such as aurora kinases [44-46], MET [47, 48], and polo-like kinases [49-51], and CRM1/XPO1 [52, 53]. The screen also pinpointed several inhibitors that differentially inhibited EMT while maintaining confluence, including inhibitors of mTOR/PI3K, DNA-PK, and p38/MAPK (**Figure 3H**). All of these pathways have been previously connected to EMT biology in prostate cancer [54-57]. We also identified the GPR40 agonist, TAK-875, and Factor Xa inhibitor, Edoxoban, as potential inducers of MET. Consistent with these observations, another GPR40 agonist, GW9508, has been shown to prevent cytokine-induced airway epithelial barriers disruption of claudin, occludin, and ZO-1 [58], and Factor Xa inhibition has been shown to reduce EMT in chronic kidney disease [59]. These agents represent promising candidates for follow-up studies to inhibit EMT and prevent or delay invasive and metastatic phenotypes associated with hormone therapy resistance.

Similar to the screen for Snail+ prostate cancer the follow-up screen for therapeutic vulnerabilities in enzalutamide-resistant CS2 cells pinpointed targets and pathways known to be involved in prostate cancer and hormone therapy resistance, including histone deacetylases, the PI3K/mTOR pathway, JAK-STAT signaling, DNA-PK, and Syk. For example, the identification of histone deacetylases and other epigenetic modifying agents is consistent with the known importance of epigenetic regulation of androgen receptor signaling [60, 61]. Other targets, however, are linked to AR signaling bypass, as in the case of PTEN loss and subsequent constitutive activation of PI3K signaling [62], activation of JAK/STAT and FGFR signaling during the acquisition of AR independence and lineage plasticity [63, 64], and the role of Syk as a potential mediator of invasive features and bone metastasis [65]. While the relevance of these targets is well supported by preclinical evidence, the clinical utility of these targets is more varied. For example, our identification of mTOR/PI3K signaling inhibition as a key vulnerability may be the result of PTEN loss in these LNCaP-derived models [66]; however, while PTEN loss is also common among patients, these agents have been unsuccessful in clinical trials [67]. Likewise, currently-available HDAC inhibitors have largely failed in clinical trials, mostly due to their toxicity [68] or lack of efficacy [69, 70]. Conversely, there are a number of ongoing clinical trials for JAK inhibitors – particularly JAK2 inhibitors – in advanced prostate cancer, but thus far these have not demonstrated sufficient monotherapy activity in men with mCRPC ([71]and see NCT00638378; closed due to lack of efficacy). Our data suggest that a number of critical and non-redundant pathways may be involved in enzalutamide resistance and lineage plasticity, suggesting the need for combination trial approaches.

Comparison across both screens identified drugs with distinct effects in a single model as well as drugs that were common hits in both screens. There are multiple possible reasons for the observed differences in hits targeting each cell line, including, but not limited to, differences in the genetic and gene expression features of each cell line [16]. For example, LNCaP95-Snail cells express AR-V7 while enzalutamide-resistant CS2 cells lack AR-V7. Enzalutamide-resistant CS2 cells also harbor dual loss of BRCA2 and RB1 and have a greater number of mutations and copy number alterations than LNCaP95 cells. These unique features may explain, at least in part, some of the differences in the list of hits from each screen.

A major limitation of the present study is the lack of *in vivo* modeling to validate the impact of our identified *in vitro* hits. This work is ongoing and also requires an assessment of the immune consequences of drug effects in the tumor microenvironment. Given the expression of mTOR and JAK/STAT signaling, for example, in immune cells and the immune suppressive impact of these agents in patients, assessing the net benefits of any drugs identified in our *in vitro* screen requires *in vivo* validation in a range of immunocompetent models either as monotherapy in selected combinations and ideally in patient correlative samples.

The current study provides a platform to quantify the effects of thousands of compounds across multiple parameters and phenotypes simultaneously to identify and prioritize candidates for follow up in a rapid and cost-effective manner. While this study is limited by the exclusive use of *in vitro* cell line models, the integration of data from phenotypic reporters, confluence imaging, and CellTiter Glo readouts across multiple models rapidly identified a prioritized list of top hits, including the dual mTOR/PI3K inhibitor, BGT-226 and the JAK2 inhibitor, AZ-960, as promising candidates for future *in vivo* studies.

## Supplemental Figure Legends

**Supplemental Figure 1. A**. Comparison of confluence for EtOH- and 4OHT-treated cells in untreated wells. **B**. Correlation between CellTiter Glo and confluence. **C**. Example of top drugs with differential growth slopes.

**Supplemental Figure 2. A**. EMT scores for isogenic pairs of enzalutamide-sensitive and - resistant cell lines using the KS scoring metric and **B**. the MLR scoring metric.

## Acknowledgments

JAS and AJA acknowledge funding support from NCI 1R01CA233585-03. MKJ was supported by Ramanujan Fellowship (SB/S2/RJN-049/2018) awarded by Science and Engineering Research Board (SERB), Department of Science and Technology, Government of India.

## Notes

### Competing Interest Statement

The authors have declared no competing interest.

