## Supplementary figures and images for "A synthetic lethal screen for Snail-induced enzalutamide resistance identifies JAK/STAT signaling as a therapeutic vulnerability in prostate cancer"

### Supplemental Figures 1-2

#### Slide 1
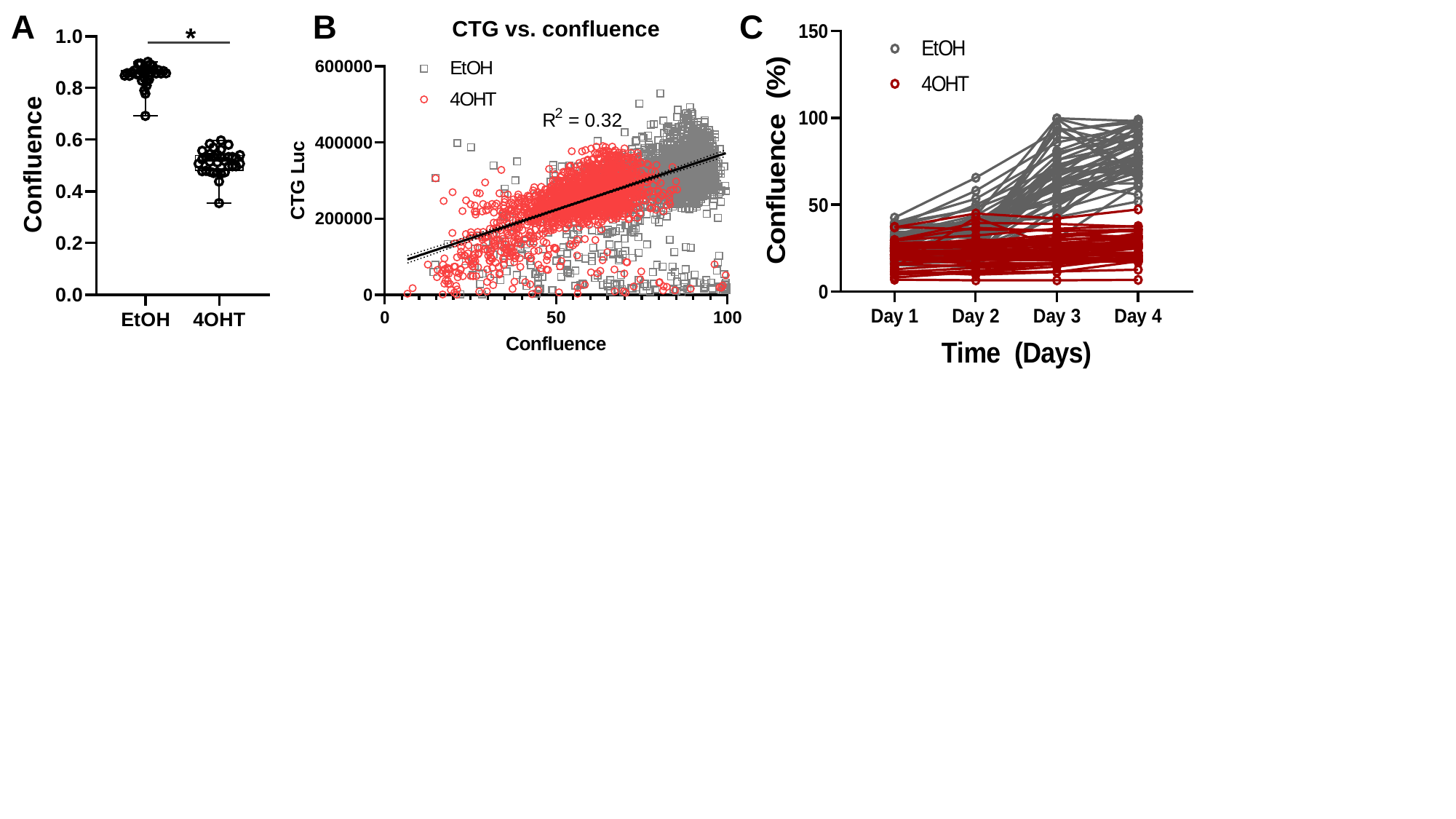

A
*
B
C

#### Slide 2
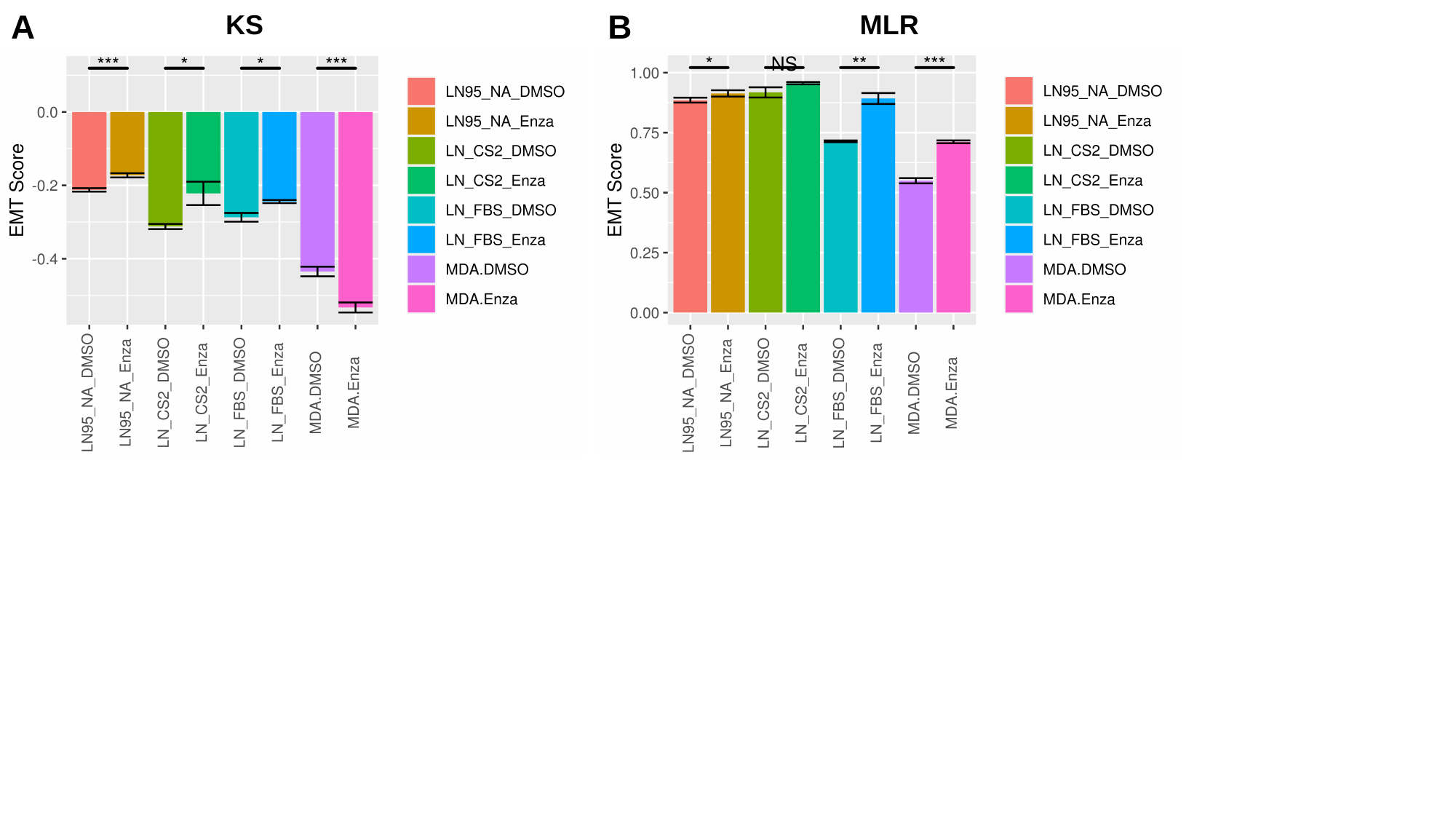

A
B
KS
MLR
